# Enhancer landscape of lung neuroendocrine tumors reveals regulatory and developmental signatures with potential theranostic implications

**DOI:** 10.1101/2023.11.17.566871

**Authors:** Ester Davis, Shani Avniel-Polak, Shahd Abu-Kamel, Israel Antman, Tsipora Saadoun, Chava Brim, Anat Bel-Ange, Karine Atlan, Tomer Tzur, Firas Abu Akar, Ori Wald, Uzi Izhar, Merav Hecht, Simona Grozinsky-Glasberg, Yotam Drier

## Abstract

Well-differentiated low-grade lung neuroendocrine tumors (lung carcinoids or LNETs) are histopathologically classified as typical and atypical LNETs, but each subtype is still heterogeneous at both the molecular level and its clinical manifestation. Here, we report the first genome-wide profiles of primary LNETs cis-regulatory elements by H3K27ac ChIP-seq with matching RNA-seq profiles. Analysis of these regulatory landscapes revealed three regulatory subtypes, independent of the typical / atypical classification. We identified unique differentiation signals that delineate each subtype. The ‘proneuronal subtype’ emerges under the influence of ASCL1, TCF4, and SOX4 transcription factors, embodying a pronounced proneuronal signature. The ‘luminal subtype’ is characterized by gain of acetylation at markers of luminal cells and GATA2 activation, and loss of LRP5 and OTP. The ‘HNF+ subtype’ is characterized by a robust enhancer landscape driven by HNF1A, HNF4A, and FOXA3, with a notable acetylation and expression of FGF signaling genes, especially FGFR3 and FGFR4 genes, pivotal components of the FGF pathway. Our findings not only deepen the understanding of LNETs’ regulatory and developmental diversity but also spotlight the HNF+ subtype’s reliance on FGFR signaling. We demonstrate that targeting this pathway with FGF inhibitors curtails tumor growth both in vitro and in xenograft models, unveiling a potential vulnerability and paving the way for targeted therapies. Overall, our work provides an important resource for studying LNETs to uncover regulatory networks, differentiation signals and therapeutically relevant dependences.

## Introduction

Lung neuroendocrine neoplasms originate from pulmonary neuroendocrine cells and account for approximately 25% of all primary lung neoplasms. They can be classified as poorly differentiated neuroendocrine carcinoma, including high-grade large cell neuroendocrine carcinoma and small cell lung carcinoma and as well-differentiated lung neuroendocrine tumors (LNETs), including typical carcinoids (grade 1), atypical carcinoids (grade 2) and carcinoids with a high mitotic/proliferation index (grade 3)^1^. However, LNETs display remarkable molecular and clinical heterogeneity and their sources are obscure. Therefore, treatment of patients with LNETs is challenging. The two main unmet needs are the lack of effective drug treatments and the lack of reliable biomarkers to guide management since the disease in patients with the same tumor grade or stage often has different clinical courses.

Specifically, there is limited data on potential molecular prognostic factors for LNETs, and there is none that have been identified to definitively and routinely guide LNETs patients’ therapeutic decisions. Previous research suggested that loss of chromosome 11q may indicate a worse prognosis upregulating anti-apoptotic pathways in LNETs^2^. Moreover, driver mutations or TP53/RB1 mutations commonly found in lung neuroendocrine carcinomas are rare in LNETs^3^ which usually retain low tumor mutational burden and PD-L1 expression^4^, limiting the possibility of using novel targeted therapies once more. Even though alterations in the mTOR pathway have not been shown to unequivocally predict treatment response, the mTORC1 inhibitor everolimus is the only FDA approved drug for unresectable LNETs, with mainly tumoristatic effect for a limited period.

The most frequently mutated family of genes in LNETs is chromatin regulators, with recurrent mutations of MEN1, ARID1A, histone methyltransferases (SETD1B, SETDB1 and NSD1) and demethylases (KDM4A, PHF8 and JMJD1C), and members of the Polycomb complex (CBX6, EZH1, and YY1)^3^. However, the genetic, epigenetic, and developmental programs that drive LNETs are still obscure, which limits our ability to identify new biomarkers and drug targets. Histone 3 lysine 27 acetylation (H3K27ac) is a histone modification that marks active enhancers^5^. In recent years, it has been demonstrated that mapping H3K27ac by chromatin immunoprecipitation sequencing (ChIP-seq) is useful for uncovering the regulatory and epigenetic state of a cell^6–8^. In particular we have recently demonstrated that pancreatic neuroendocrine tumors consist of at least two regulatory subtypes with different enhancer landscape^7^.

Here, we report on H3K27ac ChIP-seq of 23 LNETs resected from patients, and several more cell lines and xenografts, as well as matching RNA-seq data, generating the first such resource. We perform elaborate analysis of the data, revealing regulatory subtypes, highlighting super-enhancers and transcription factors that drive these regulatory programs, and uncovering subtype-specific dependency on FGFR signaling that can suggest new potential therapeutic avenues.

## Results

### Genome-wide profiles of LNETs cis-regulatory elements distinguishes three regulatory subtypes

We profiled the H3K27ac landscape of 16 primary typical and 6 primary atypical lung neuroendocrine tumors, and one skin metastasis of atypical lung neuroendocrine tumor (Suppl. Table 1). We identified 25,372 peaks overlapping known promoters, and 44,140 which do not, representing putative enhancers.

**Table 1:** Top ten MsigDB terms enriched in genes near enhancers significantly more acetylated (fold-change > 2, FDR < 10%) in proneural LNETs compared to HNF+ LNETs.

| **MSigDB Term** | **p-value** | **Relative Enrichment** |
| --- | --- | --- |
| MEISSNER_BRAIN_HCP_WITH_H3K4ME3_AND_H3K27ME3 | 3.62E-10 | 1.80 |
| GO_SYNAPSE | 2.86E-09 | 1.90 |
| ACEVEDO_METHYLATED_IN_LIVER_CANCER_DN | 6.46E-09 | 1.97 |
| GO_SYNAPSE_PART | 8.37E-09 | 1.99 |
| BENPORATH_ES_WITH_H3K27ME3 | 1.60E-08 | 1.77 |
| MIKKELSEN_MEF_HCP_WITH_H3K27ME3 | 3.30E-08 | 2.08 |
| GO_CELL_CELL_SIGNALING | 3.57E-08 | 1.90 |
| GO_GATED_CHANNEL_ACTIVITY | 4.94E-08 | 2.39 |
| GO_POSTSYNAPSE | 9.43E-08 | 2.20 |
| BENPORATH_SUZ12_TARGETS | 9.96E-08 | 1.76 |

We compared the global enhancer landscapes of profiled LNETs to previously published enhancer landscapes of ileal and pancreatic neuroendocrine tumors^7^. 84% (21,242) of the LNET promoter H3K27ac peaks are shared with other NETs, compared to only 65% (28,658) of the putative enhancers. Shared peaks across neuroendocrine tumors are enriched in motifs of AP1, FOX, RFX, IRX, HNF, ETS and others (Suppl. Table 2). Comparing acetylation levels across neuroendocrine tumors revealed 3,866 peaks that are significantly more acetylated (DESeq2, fold-change > 2, FDR < 10%, see methods) in LNETs and 3,231 that are more acetylated in ileal and pancreatic neuroendocrine tumors. LNET specific enhancers were enriched with the different motifs, including snail/slug, ZSCAN, SOX, T-box and ASCL (Suppl. Table 2). We clustered all samples based on the pairwise Spearman correlations of their enhancer landscape. The heterogeneity of LNETs enhancer landscape is higher than that of ileal neuroendocrine tumors, and even higher than that of pancreatic neuroendocrine tumors, as evident by the low correlation between different LNET clusters (Fig. 1a).

**Table 2:** Top ten MSigDB terms enriched in genes near enhancers significantly more acetylated (fold-change > 2, FDR < 10%) in HNF+ LNETs compared to proneural LNETs.

| MSigDB Term | p-value | Relative Enrichment |
| --- | --- | --- |
| SERVITJA_ISLET_HNF1A_TARGETS_DN | 9.44E-13 | 4.59 |
| MEISSNER_BRAIN_HCP_WITH_H3K4ME3_AND_H3K27ME3 | 2.15E-10 | 1.81 |
| LI_WILMS_TUMOR_VS_FETAL_KIDNEY_1_UP | 1.47E-08 | 2.88 |
| BENPORATH_ES_WITH_H3K27ME3 | 1.75E-08 | 1.76 |
| BENPORATH_SUZ12_TARGETS | 5.05E-08 | 1.77 |
| GO_REGULATION_OF_LIPID_METABOLIC_PROCESS | 5.83E-08 | 2.49 |
| ONDER_CDH1_TARGETS_2_DN | 1.33E-07 | 2.10 |
| HSIAO_LIVER_SPECIFIC_GENES | 3.07E-07 | 2.56 |
| FLECHNER_KIDNEY_TRANSPLANT_REJECTED_VS_OK_DN | 1.00E-06 | 1.85 |
| CHYLA_CBFA2T3_TARGETS_DN | 1.20E-06 | 2.46 |

**Figure 1:**
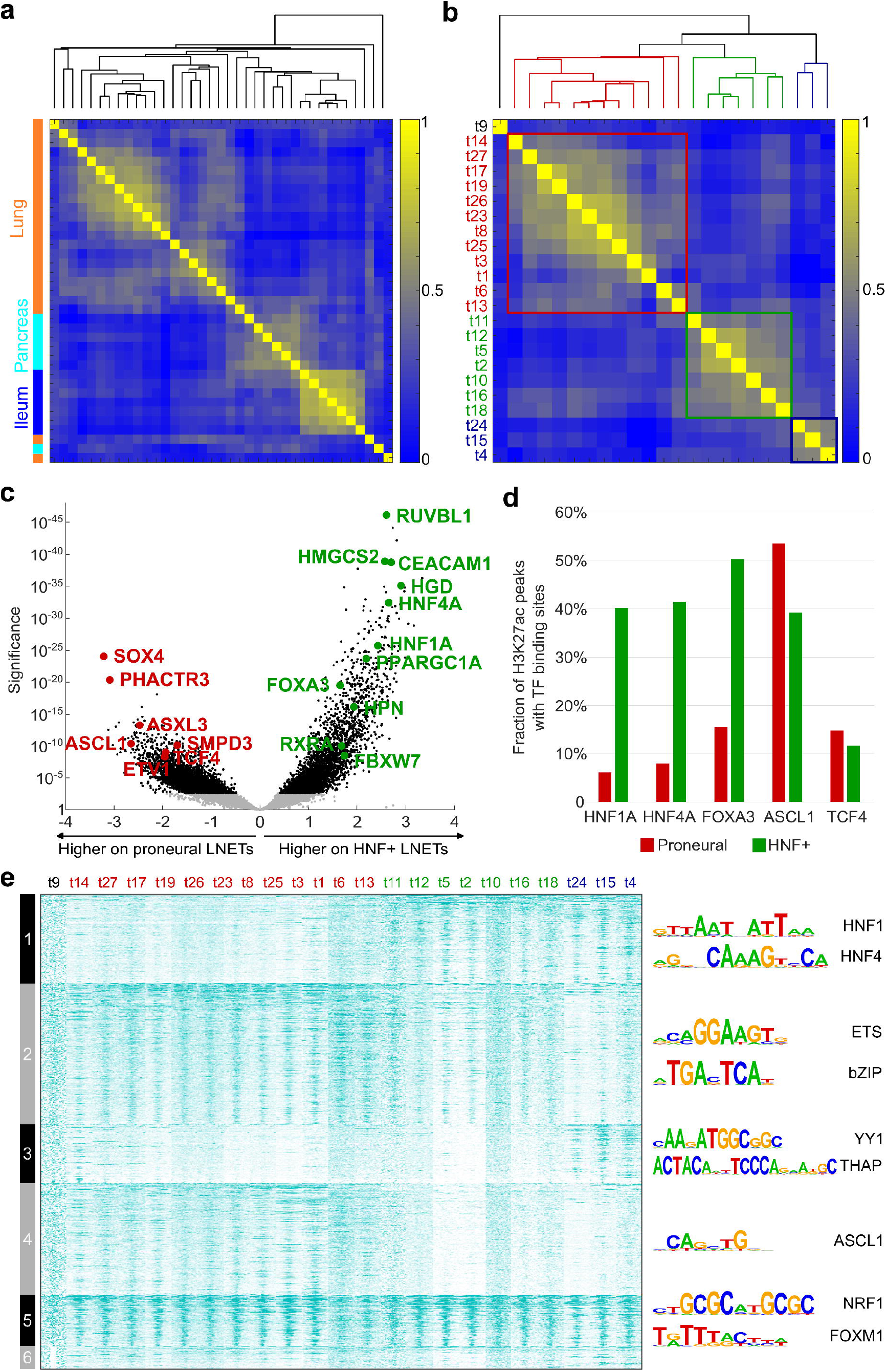
Regulatory and developmental subtypes of lung neuroendocrine tumors. **(a)** Pairwise Spearman correlations of H3K27ac signals at distal regulatory elements across lung, pancreatic and ileal neuroendocrine tumors demonstrate high heterogeneity between lung NETs clusters **(b)** Pairwise Spearman correlations of H3K27ac signals at distal regulatory elements in 23 lung neuroendocrine tumors suggest three possible regulatory subtypes. Tumors in the first cluster (termed proneural LNETs) are marked in red, second cluster (termed HNF+ LNETs) marked in green, and the third cluster (termed luminal LNETs) in blue. **(c)** Significance (log_10_ scale, Wald’s test) and log_2_ fold differences in H3K27ac ChIP-seq signals in HNF+ and proneural LNETs, calculated by DESeq2 comparison of 19 biologically independent tumors. Each dot represents an individual site (black: FDR□:<□:0.01). Selected H3K27ac peaks are marked by red circle (higher in proneural) or green circle (higher in HNF+) and annotated by the nearest gene. **(d)** Percentage of proneural and HNF+ specific H3K27ac peaks with binding sites for HNF1A, HNF4A, FOXA3, ASCL1 and TCF4. HNF+ specific peaks are strongly enriched with HNF1A, HNF4A and FOXA3 binding sites. **(e)** Heat map showing normalized acetylation levels at the union set of enhancers across all LNET samples. Enhancers are clustered by k-means clustering, bars on the left represent the 6 clusters. Sample names are on top, proneural LNETs are colored red, HNF+ LNETs in green and luminal LNETs in blue. T9 is an outlier that could not be classified, potentially due to lower ChIP quality. Top enriched motifs in each cluster are shown on the right. Enhancer cluster 5 is common to all tumors. Proneural tumors are also positive for clusters 2 and 4, HNF+ for clusters 1 and 2 and luminal tumors for clusters 1 and 3.

Next, we focused only on LNETs, to better classify the tumors. We quantified acetylation at all LNET H3K27ac peaks not containing a TSS (putative enhancers) and performed hierarchical clustering based on Spearman correlation. This uncovered three clusters (Fig. 1b), in agreement with previous integrative analyses^9,10^. One cluster contained only three tumors, and two major clusters contained 7 and 12 tumors. The subtypes were independent of the typical/atypical classifications, as each of the clusters contains both typical and atypical tumors. The only sample taken from a metastatic site (t14) also did not differ significantly for the rest and seemed to match its cluster.

### Characterization of the unique regulatory landscape of each of the three LNETs subtypes

To better understand the difference between the clusters we identified the H3K27ac peaks that are more acetylated in each cluster and the genes near them. Proneural transcription factors and their targets were found enriched near peaks specific to the first cluster, and we have therefore termed this subtype “proneural”. In particular, these include the enrichment of multiple synapse related GO terms (Table 1, Suppl. Table 3). These tumors exhibit strong acetylation near related genes, such as ASCL1, SOX4, TCF4, SMPD3, PHACTR3, ASXL3, and ETV1 (Fig. 1c). Proneural-specific peaks were enriched with binding motifs of ASCL1, SOX4, TCF4, ETV1, Snail/Slug, ZEB, and other related transcription factors, suggesting that they bind to enhancers and regulate transcription in proneural NETs (Suppl. Table 2).

The second subtype exhibited specific acetylation near HNF targets, and we therefore termed it “the HNF+ subtype”. In particular, we observed enrichment of HNF1A targets, liver-specific genes and genes involved in lipid metabolism (Table 2, Suppl. Table 3). Indeed, these tumors exhibited strongly acetylated enhancers near related genes such as HNF1A, HNF4A, CEACAM1, HMGCS2, FOXA3, PPARGC1A, RXRA, FBXW7, HGD, and HPN (Fig. 1c, Supp. Fig. 1a-e). Motif analysis of HNF+ LNETs H3K27ac peaks revealed strong enrichment of HNF1, HNF4, FOXA, and PPAR/RXR/TR motifs (Suppl. Table 2). Notably, the H3K27ac peak at the HNF1A locus spans the promoters of both HNF1A and a lncRNA known as HNF1A-AS1 transcribed from the other strand. Both genes are expressed specifically in HNF+ LNETs, and HNF1A-AS1 to an even greater extent (Fig. 2a).

**Figure 2:**
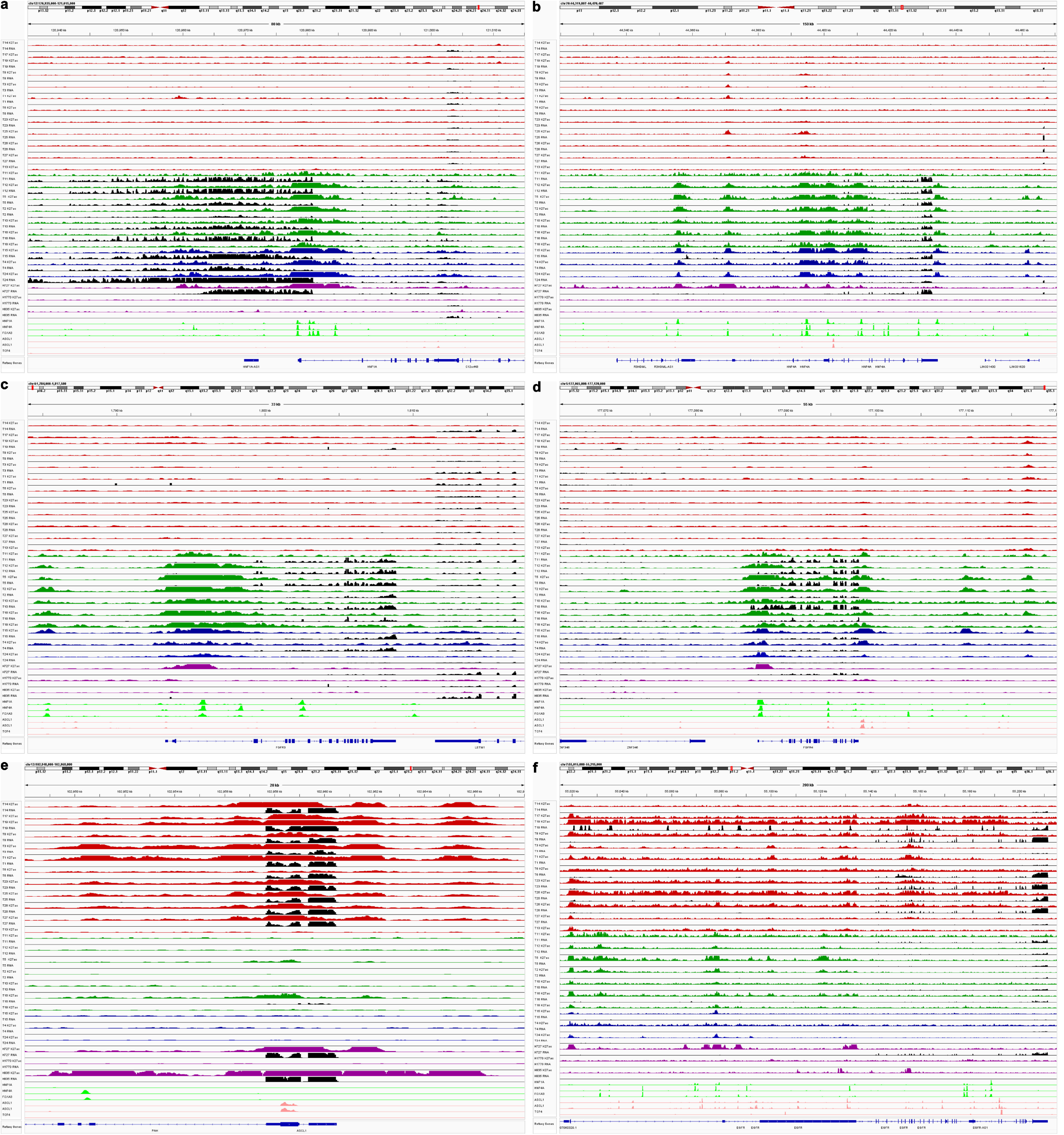
HNF1A, HNF4A, FGFR3 and FGFR4 demonstrate strong enhancer acetylation and expression specifically at HNF+ LNETs, while the ASCL1 and EGFR locus is preferentially acetylated in proneural LNETs. **(a-e)** Genomic view of the **(a)** HNF1A, **(b)** HNF4A, **(c)** FGFR3, **(d)** FGFR4, **(e)** ASCL1, and **(e)** EGFR loci, showing RNA signal (black) and H3K27ac signals of proneural (red), HNF+ (green) and luminal (blue) lung neuroendocrine tumors, as well as the NCI-H727, NCI-H1770 and NCI-H835 lung neuroendocrine neoplasm cell lines (purple). Hepatic factors (HNF1A, HNF4A and FOXA3) ChIP-seq tracks in HepG2 cells are shown in light green and proneural transcription factors (ASCL1 and TCF4) ChIP-seq tracks in neuroblastoma cells in light red. RNA-seq signals are normalized to 5 bins per million mapped reads (BPM). H3K27ac signals are scaled by promoter-based DESeq2 normalization (see Methods). Multiple HNF1, HNF4 and FOXA3 binding sites are found at FGFR3 and FGFR4 enhancers and promoters, suggesting FGFR expression is downstream of hepatic factors activation.

The third smaller subtype was enriched in acetylated genes related to luminal identity and is therefore termed here “luminal subtype”. This is reflected in the enrichment (Suppl. Table 3) of genes down-regulated in basal subtype of breast cancer (SMID_BREAST_CANCER_BASAL_DN, p<1.1*10^−10^), genes up-regulated in luminal-like breast cancer cell lines compared to the basal-like ones (CHARAFE_BREAST_CANCER_LUMINAL_VS_BASAL_UP, p<2*10^−9^) and genes down-regulated in ER-TMX2-28 breast cells compared to luminal MCF7 cells (GOZGIT_ESR1_TARGETS_DN, p<4*10^−8^). Moreover, when considering H3K27ac peaks that lose acetylation in this subtype (i.e., are more acetylated in the other LNET tumors), we detect enrichment for genes up-regulated in mammary stem cells (LIM_MAMMARY_STEM_CELL_UP, p<9*10^−10^) and genes down-regulated in mature mammary luminal cells (LIM_MAMMARY_LUMINAL_MATURE_DN, p<5*10^−7^). The luminal subtype is more similar to the HNF+ subtype, with strong acetylation at HNF1A, HNF4A, CEACAM1, HMGCS2, HGD, HPN, and FOXA3 (Fig. 2a,b, Supp. Fig. 1a-e), and with overall higher correlation of the genome-wide enhancer landscape (Fig. 1b). However, it lacks FBXW7 acetylation and does exhibit SOX4 acetylation (Supp. Fig. 1f), one of the key markers of proneural LNETs. It also displays unique strong enhancers near several genes such as FEV, GATA2, TSSC4, CCKBR, RET, CCNP-AKT2, ANGEL1, and PARVB (Supp. Fig. 1i-j). The two most enriched motifs in H3K27ac peaks unique to luminal LNETs are the FOXA (p<10^−120^) and GATA2 (p<10^−119^) motifs (Suppl. Table 2), in line with these results. Importantly, GATA is required for the luminal identity of mammary cells^11,12^.

We clustered the enhancers to six clusters based on their acetylation across the samples (Fig. 1e), showing robust acetylation across the different clusters. Cluster 1, shared by HNF+ and luminal LNETs is enriched for HNF1 and HNF4 motifs, consistent with the shared acetylation profiles of HNF+ and luminal LNETs. Cluster 2, shared by proneural and HNF+ LNETs is enriched for ETS and bZIP motifs. Proneural LNETs showed higher ETS activity than HNF+ LNETs and were ETV1 positive (Supp. Fig. 1h), but the luminal specific loss of ETS enhancers may be due to activation of FEV (Supp. Fig. 1j) that binds ETS binding sites and repress them^13^. Cluster 3, unique to luminal LNETs, is enriched with YY1 and THAP11 motifs. Cluster 4, specific for proneural LNETs, is enriched with ASCL1 motif, in line with the findings above. Cluster 5 enhancers are common to all tumors and are enriched with NRF1 and FOXM1 motifs. The enhancers in cluster 6 demonstrate only a low level of acetylation, and are not enriched in any known motif. The fact that proneural LNETs are positive for clusters 2 and 4, HNF+ for clusters 1 and 2 and luminal tumors for clusters 1 and 3, suggests the proneural and luminal subtypes present the two ends of a regulatory spectrum and HNF+ LNETs contain some of the features of the both the proneural and luminal subtypes.

### Analysis of transcription factor binding within enhancers supports differential developmental trajectories

Next, we tested whether the sequence motifs that we identified in the differential enhancers represent actual differential binding sites of ASCL1, TCF4, HNF1A, and HNF4A and FOXA3 transcription factors. We utilized ASCL1^14^ and TCF4^15^ ChIP-seq data from neuroblastoma cells, and HNF1A, HNF4A, and FOXA3 ChIP-seq data from HepG2 cells (data from ENCODE^15^).

A significant portion of 53.4% (1610 / 3016) of proneural specific peaks overlapped with ASCL1 binding sites (p<3*10^−124^, Fisher exact test), but only 39.2% (855 / 2181) of HNF+ peaks displayed ASCL1 binding. 14.7% (444 / 3016) of the proneural peaks overlapped the TCF4 binding sites (p<5*10^−35^, Fisher exact test), but only 11.6% (252 / 2181) of the HNF+ peaks displayed TCF4 binding. More strikingly, 40.1% (875 / 2181, Fisher exact test p<10^−167^), 41.4% (903 / 2181, Fisher exact test p<6*10^−251^) and 50.2% (1095 / 2181, Fisher exact test p<4*10^−153^) of HNF+ LNET peaks demonstrated HNF1A, HNF4A, and FOXA3 binding, respectively, compared to only 6.1% (184 / 3016), 7.9% (239 / 3016) and 15.4% (465 / 3016) of proneural peaks (Fig. 1d). This suggests that many of the differential enhancer landscapes are due to HNF1A, HNF4A, and FOXA3 activation in HNF+ LNETs, and possibly due to ASCL1 and TCF4 activation in proneural LNETs. Indeed, many liver-specific genes have HNF+ LNET specific peaks bound by HNF1A and HNF4A, including genes involved in drug metabolism, such as CYP4F3 and CYP3A5, as well as the HNF4A and HNF1A genes themselves (Fig. 2a,b). Although CEACAM1 is not a liver-specific gene, it has been identified as a target of HNF1A^16^ and HNF4A^17^, and its binding can be detected in an HNF+ specific enhancer upstream of CEACAM1 (Supp. Fig. 1a). Importantly, CEACAM1 facilitates metastasis in multiple cancers^18–20^, suggesting it may contribute to the metastatic process of HNF+ LNETs. Given that the main distant metastatic site of LNETs is the liver, it may suggest the expression of HNFs and their targets may also contribute to liver colonization.

These results validate the binding of ASCL, HNF1A, and HNF4A transcription factors to differential enhancers. It suggests that many of the HNF+ LNETs differential enhancer landscapes are due to HNF1A, HNF4A, and FOXA3 activation, and ASCL1 and TCF4 activation in proneural LNETs.

### Super-enhancers are differentially acetylated between LNET subtypes

Next, we used the H3K27ac ChIP-seq to identify super-enhancers. Across all tumors 2254 super-enhancers were identified. Of those, 561 overlapped with the proneural-enriched enhancers, 545 with HNF+ enriched enhancers, and another 399 were enriched only in the luminal subtype, demonstrating that most super-enhancers (67%) were differentially acetylated between the subtypes. Many of the enhancers mentioned above are in fact super-enhancers such as ASCL1, SOX4, TCF4, SMPD3, PHACTR3, ASXL3, ETV1, HNF1A, HNF4A, CEACAM1, PPARGC1A, FOXA3, RXRA, FBXW7, HGD, TSSC4, FEV, GATA2, CCKBR, CCNP-AKT2, ANGEL1, and PARVB.

The 20 most differentially acetylated super-enhancers between the luminal LNETs and the other LNETs were all more acetylated in luminal LNETs except two loci-LRP5 and OTP. These luminal specific super-enhancers include the loci mentioned above TSSC4, CCKBR, CCNP-AKT2, and ANGEL1. The 20 most differentially acetylated super-enhancers between the proneural and HNF+ LNETs were all more strongly acetylated in HNF+ LNETs, and included many interesting loci such as RUVBL1 (#1), HGD (#4), PDGFA (#6), TM4SF5 (#8), IL1RL2 (#11), and FGFR3 (#12).

### A subtype-specific dependency of HNF+ LNETs on FGFR signaling

Of the above super-enhancers, we decided to focus on FGFR3 since it is important for tumor growth and can be targeted by small-molecule drugs. Examination of other FGFR family members revealed that FGFR4 is also strongly acetylated specifically in HNF+ LNETs. Interestingly, the HNF+-specific enhancers at FGFR3 and FGFR4 are bound by HNF1A and HNF4A, suggesting that they support the acetylation and expression of FGFRs (Fig. 2c,d).

To identify alternative receptor tyrosine kinase signaling in the proneural subtype, we examined proneural super-enhancers near RTK-related genes. These included EGFR, HGF, and PDGFB, the most prominent of which was EGFR (Fig. 2f). EGF signaling has been previously suggested to drive LNETs growth and its inhibition as an effective therapy^21,22^. Based on our analysis, we hypothesized that this is specifically true for proneural LNETs, but less so for HGF+ LNETs, emphasizing the importance of this classification for therapeutic applications.

Therefore, we tested FGF inhibition by treating three human lung neuroendocrine neoplasm cell lines-two of well-differentiated LNET origin (NCI-H727 and NCI-H835), and one of the lung neuroendocrine carcinoma origin (NCI-H1770) with either Erdafitinib or Lenvatinib for 24, 48, and 72 h, and tested their impact on cell viability. Lenvatinib is a multi-targeting tyrosine kinase inhibitor that targets the VEGFR and FGFR receptors and has been reported to be effective in treating advanced pancreatic and gastrointestinal NETs^23^. Erdafitinib is a pan-FGFR tyrosine kinase inhibitor, with minimal activity against VEGFR kinases.

Of the three, only NCI-H727 cells showed sensitivity to Lenvatinib and Erdafitinib (Fig. 3a), with less than 50% viable cells at 15μM, suggesting that this is the only cell line that relies on FGF signaling. In contrast, NCI-H835 cells demonstrated remarkable resistance to both Lenvatinib and Erdafitinib. We profiled enhancers by H3K27ac ChIP-seq in these cell lines and, as expected, only NCI-H727 had an active enhancer at the FGFR3 locus (Fig. 2c). The global enhancer landscape of the cell lines was considerably different from that of the primary LNETs (Supp. Fig. 2), probably due to the inherent limitations of in vitro cell line models, such as different environmental factors, in vitro selection pressures, and genetic drift and instability. However, NCI-H727 cells retained the main HNF+ marker enhancers near HNF1A, HNF4A, CEACAM1, FOXA3, HGD, and HPN (Fig. 2a-b, Supp. Fig. 1a,c-e). NCI-H835, the line most resistant to FGF inhibition, displays acetylation of the key proneural enhancers ASCL1, SMPD3, SOX4, and ETV1 (Fig. 2e, Supp. Fig. 1f-h), although some were also positive for NCI-H727.

**Figure 3:**
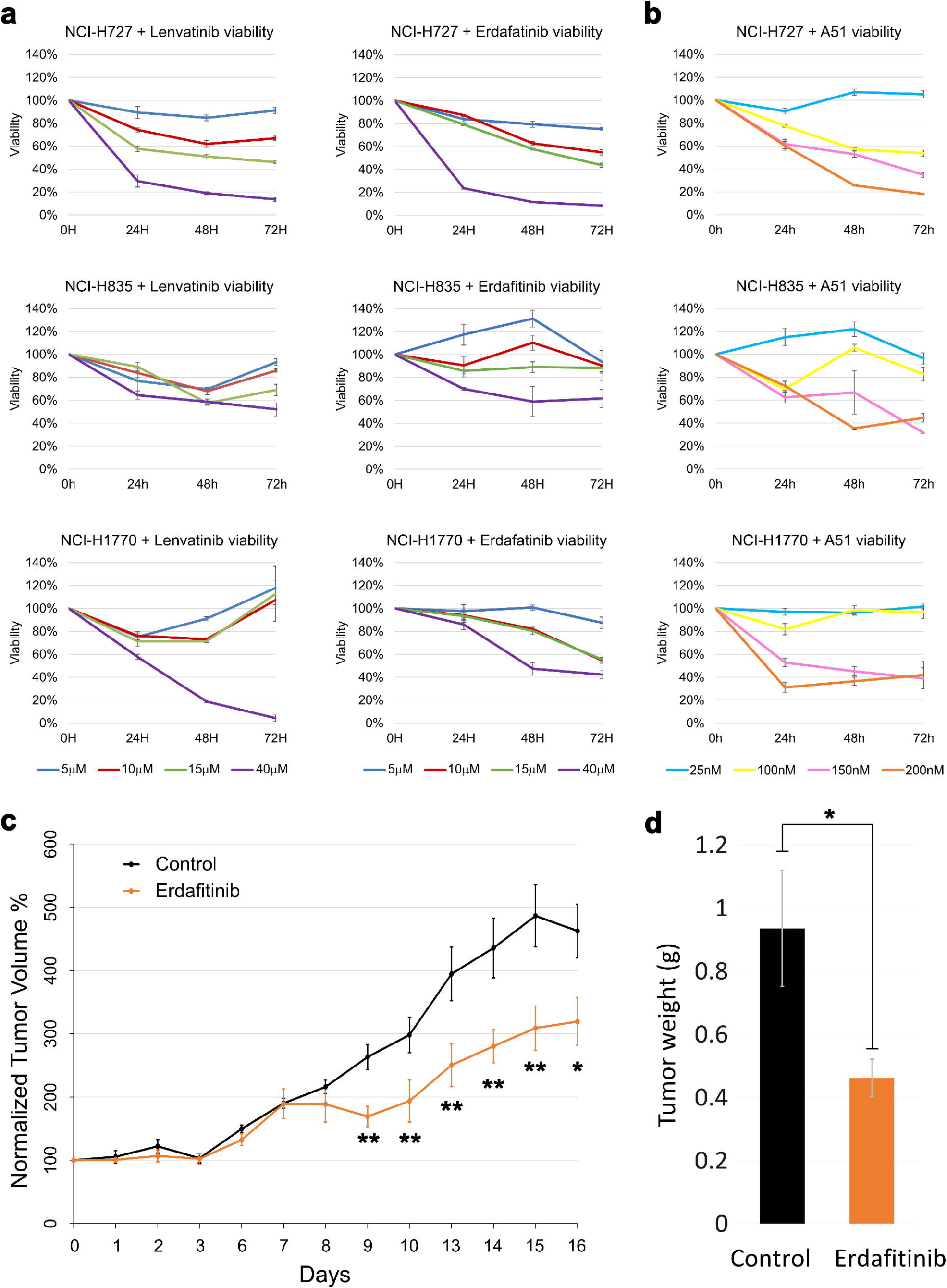
Erdafitinib significantly slows tumor growth. **(a)** Relative cell line viability as measured by WST-1 assay of NCI-H727, NCI-H835 and NCI-H1770 lines during treatment with Lenvatinib (left) or Erdafitinib (right). Values represent the ratio between viability of treated cells and of control cells treated with DMSO. Error bars represent standard error of the mean. **(b)** Relative cell line viability as measured by WST-1 assay of the NCI-H727, NCI-H835 and NCI-H1770 during treatment with A51. Values represent the ratio between viability of treated cells and of control cells treated with DMSO. Error bars represent standard error of the mean. **(c)** Normalized tumor volume of NCI-H727 mouse xenograft model treated with Erdafitinib (12.5mg/kg, orange) or PBS 1% tween control (black). Error bars represent standard error of the mean. P-values are calculated by one tail t-test, * p < 0.05, ** p < 0.01. **(d)** Average tumor weight of the PBS control group (black) and Erdafitinib treated group (orange) at the end of the experiment after mice were sacrificed. Error bars represent standard error of the mean. P-values are calculated by one tail t-test, * p < 0.05.

Taken together, these results suggest that HNF+ NETs are sensitive to FGFR inhibitors, while other subtypes may be more resistant.

### LNET models show sensitivity to super-enhancer inhibition by A51

Since NCI-H835 is not sensitive to FGFR inhibition, but still display strong acetylation at many proneural NET super-enhancers, we next tested whether it may be sensitive to the inhibition of super-enhancer activity by the novel small-molecule multi-kinase inhibitor A51^24^. All three cell lines showed similar sensitivities to A51 (Fig. 3b), suggesting that super-enhancer-targeting drugs may be helpful across lung neuroendocrine subtypes. At the molecular level, the effect of A51 was cell line-specific, where genes downregulated by A51 were enriched in the Neuron Development and Synapse GO terms in NCI-H835 cells and in liver-specific genes in NCI-H727 cells (Suppl. Table 3).

These results suggest that super-enhancers drive the proneural and HNF+ patterns of NCI-H835 and NCI-H727, respectively, and that A51 can target this identity by perturbing super-enhancer function. Interestingly, liver-specific genes were upregulated by A51 in NCI-H835 cells, suggesting that once neural identity is lost, HNF+ differentiation is activated.

### A mouse xenograft HNF+ LNET model is sensitive to FGFR inhibition

Erdafitinib demonstrated high efficacy in NCI-H727 cells, and may therefore be a promising new direction for the treatment of HNF+ LNETs. To further test this hypothesis in vivo, we generated a mouse xenograft LNET model by subcutaneous injection of NCI-H727 cells into the back of athymic nude mice. The H3K27ac profiles of tumors obtained from this model were very similar to those of NCI-H727 cells grown in vitro (Supp. Fig. 2), suggesting that the differences observed between cell lines and primary tumors are not due to differences in the microenvironment or environmental factors and cannot be easily reversed. Importantly, daily treatment of these mice with Erdafitinib (12.5 mg/kg) significantly slowed tumor growth in vivo (Fig. 3c). Repeat of this experiment yielded similar results (Supp. Fig. 3), demonstrating its robustness. Treatment with Lenvatinib (5 mg/kg) may also slow tumor growth, but to a somewhat lesser extent (Supp. Fig. 3b,c).

## Discussion

We report on three LNET subtypes based on their genome-wide enhancer profiles. Comprehensive analysis of their enhancer landscape uncovered regulatory and differentiation signatures of each subtype. One subype is characterized by activation of proneural transcription factors, another by hepatic nuclear factors, and the third displaying luminal-like features. The last two subtypes, and especially the HNF subtype, show strong acetylation at a super-enhancer in the FGFR3 locus, suggesting a dependency on FGF signaling. Indeed, we show FGF inhibition slows down tumor growth both in vitro and in vivo, demonstrating the utility of enhancer profiles to uncover new dependencies and therapeutic avenues.

Our study raises important hypotheses about the origin and development of different LNETs subtypes. Of particular interest is the HNF+ subtype which resembles hepatic differentiation. Pulmonary neuroendocrine cells and hepatocytes both differentiate from the endoderm, with evidence of de-differentiation or transdifferentiation between hepatic and neuroendocrine cells. For example, induced transdifferentiation^25,26^ and the occurrence of hepatocellular carcinoma with neuroendocrine differentiation and mixed hepatocellular carcinoma-neuroendocrine carcinoma tumors^27–29^. As mentioned above (Table 1), proneural-specific enhancers are enriched in genes methylated in the normal liver (and hypomethylated in hepatocellular carcinoma), consistent with the notion of the hepatic-neuroendocrine transdifferentiation axis, and that the HNF+ LNETs are closer to the hepatic end of this axis. Interestingly both the proneural and the HNF+ subtypes demonstrated strong enrichment for acetylation near bivalent high CpG density promoters in the brain (Tables 1 and 2). 214 of the 220 enriched genes had acetylated peaks near them in only one of the subtypes, suggesting these bivalent genes become differentially active in each subtype, further supporting the notion that we are able to capture differentiation signals of relevant genes in the different subtypes.

The luminal subtypes exhibit GATA2 super-enhancer and the GATA motif is enriched in luminal specific H3K27ac peaks. Since GATA3 is required for the luminal lineage identity of mammary cells^11,12^ it raises the hypothesis that GATA2 contributes to the luminal identity of luminal LNETs. Moreover, the LRP5 super-enhancer common to other LNETs but entirely missing in luminal LNETs is of interest (Supp. Fig. 1k). The Wnt receptor LRP5 is required for mammary ductal stem cells^30^ and to maintain mammary basal lineage^31^, suggesting loss of the LRP5 super-enhancer may drive luminal LNETs towards luminal identity. The loss of the Orthopedia homeobox protein (OTP) super-enhancer may also be involved (Supp. Fig. 1l), since OTP controls the differentiation of hypothalamic neuroendocrine cells^32,33^. It is also important clinically since OTP was suggested as a biomarker for LNETs^34,35^, but our findings imply it will fail to identify luminal LNETs. The enhancer cluster unique to the luminal LNETs (Cluster 3) is enriched with YY1 and THAP11 motifs. THAP11 is a transcriptional repressor, and the THAP domain is known to promote transcriptional repression^36^. YY1 is a transcriptional partner of THAP1 and loss of THAP1 also disrupts YY1 binding^37^. THAP12 expression is lower in luminal LNETs, and therefore if THAP12 shares a similar function to THAP1 and THAP11, its loss may explain the acetylation gain at THAP11 and YY1 binding sites.

Several findings have clear clinical significance and can serve as a basis for future clinical studies: 1) LNET patients may benefit from super-enhancer targeting drugs such as A51; 2) Patients with HNF+ LNETs may benefit from FGFR inhibition-based treatment; 3) Specific FGFR inhibitors such as Erdafitinib may be a better fit than broader kinase inhibitors such as Lenvatinib; and 4) personalized treatment based on subtype can help fit the right drug to the right patient, and subtypes can be identified by a few biomarkers highlighted in this study. Overall, this study demonstrates the utility of regulatory profiling of tumors by H3K27ac, provides an important resource for the community, and raise a few clinically relevant hypotheses and novel therapeutic avenues.

## Methods

### Clinical materials

Fresh-frozen LNET specimens were obtained from patients treated at the Neuroendocrine Tumor Unit, ENETS Center of Excellence, Hadassah Medical Center, Jerusalem, Israel, under the Hadassah Institutional Helsinki committee, approval no. 0627-20-HMO.

### RNA-seq

2 mm tissue sample was homogenized in tri-reagent, and RNA was extracted using the standard tri-reagent protocol. After precipitation, the RNA was resuspended in an appropriate volume of RNase-free water supplemented with RNA inhibitors (New England Biolabs). RNA samples were stored at -80°C until further use. The quality of the extracted RNA was assessed using the Agilent TapeStation System. 250 ng RNA was used to generate RNA-seq libraries using the KAPA mRNA Hyperprep Kit, according to the manufacturer’s instructions.

### Chromatin immunoprecipitation sequencing (Chip-seq)

Chip-seq was performed as previously described^38^. Briefly, cultured cells or minced frozen tissues were cross-linked in 1% formaldehyde. Sonication of the samples was calibrated such that DNA was sheared to 300-700bp fragments. Histone H3K27 acetylation was immunoprecipitated using an antibody from the Active Motif (no. 39133). ChIP DNA was used to generate sequencing libraries using a KAPA Hyper Prep Kit.

### Cell lines and inhibitors experiments

Lenvatinib and Erdafitinib (JNJ-42756493) were purchased from MedChemExpress (NJ, USA). The A51 inhibitor was a gift from Prof. Yinon Ben-Neriah. NCI-H727, NCI-H1770, and NCI-H835 cell lines were acquired from the ATCC. The cell lines were cultured in complete RPMI 1640 medium (Gibco, Thermo Fisher Scientific, Massachusetts, USA) supplemented with 10% or 5% fetal bovine serum (FBS; HyClone, Utah, UK), 1% penicillin streptomycin (Diagnovum, Tilburg, Netherlands), 1% L-glutamine (Sartorius, Göttingen, Germany), 1% sodium pyruvate (Sartorius), and 1% non-essential amino acids (Diagnovum). The WST-1 cell viability assay (Abcam, Cambridge, England) was performed on the three cell lines by seeding cells in 100μl into individual wells of a 96-well plate. Cells were treated with different concentrations of inhibitors or DMSO control for 24h, 48h and 72h. After the respective incubation periods, 10μl of WST-1 reagent was added to each well and incubated for 2-4 h. The plates were then scanned using a microplate spectrophotometer (Multiskan GO, Thermo Scientific, Massachusetts, USA) at wavelengths of 450 and 650 nm. The resulting data were analyzed to assess the cell viability at each concentration and time point.

### In-vivo inhibitor experiments

LNET xenografts were established 5 days after the subcutaneous injection of 4 × 10^6^ NCI-H727 cells into the back of athymic nude mice (FOXN1NU NU/NU mice). Once the neoplasm size reached 130mm^3^, the mice were randomized into two groups and treated for the next 4 weeks with (1) vehicle1% Tween 80 and (2) erdafitinib (JNJ-42756493) 12.5 mg/kg dissolved in 1% Tween 80 (N=7). Animals were treated daily (except on weekends) for 20 days, and the drugs were administered by oral gavage. Tumor size was measured daily using a caliper, and tumor volume was estimated using the following equation: length × (width^2^)/2. Drug concentrations were chosen after conducting calibration experiments at the recommended concentrations. The experiments were repeated using the same model, with minor changes. Mice were randomized into three groups and treated for the next 4 weeks with: (1) vehicle 0.5% carboxymethylcellulose (CMC), (2) lenvatinib (5 mg/kg) dissolved in 0.5% CMC, and (3) erdafitinib (JNJ-42756493) (12.5 mg) dissolved in 1% Tween 80. The animals were treated daily (N=6) and the tumor volume was calculated as described previously.

### Computational and statistical analyses

ChIP-seq reads were aligned to the reference genome (hg38) using the BWA 0.7.17 (Ref. ^39^). Reads with a mapping quality < 10 were not counted, and reads aligned to the same position and strand were counted only once. Density signals were calculated using the deeptools bamCoverage^40^ and visualized on the Integrated Genome Viewer^44^. Regular H3K27ac peaks were called using HOMER^41^, with style ‘histone’ for variable length, and super-enhancer using style ‘super’ and requiring a local fold change of 1 (-L 1). We assigned enhancers to the nearest promoter for differential super-enhancers, with multiple nearby genes assigned to the nearest gene, which was more highly expressed in the matching subtype.

We generated a union set of H3K27ac peaks by taking the top 40,000 peaks of each sample, removing ENCODE blacklisted peaks, and merging all remaining peaks using bedtools merge^42^. Acetylation signal was estimated from ChIP-seq reads that fell within this union set of H3K27ac peaks. To compare differential acetylation at peaks between samples, we used DESeq2 (Ref. ^43^) for read counts computed using featureCounts^44^. Promoter signals (<2 kb downstream and <2.5 kb upstream of TSSs) vary less than those at enhancers and were therefore used to calculate normalization factors (size factor, SFs) for each library by DESeq2. When comparing libraries normalized by these SF, we considered only enhancers with average normalized read counts ≥ 50. *P* values were calculated using the Wald test and false discovery rates (FDRs) using the Benjamini-Hochberg method^45^. Enhancers with a normalized read count > 2-fold and FDR < 0.01 were considered enriched in the given subtype. Since the cohort contained only 3 luminal LNETs, we first compared proneural against HNF+ LNETs. Afterwards, luminal LNETs were compared against proneural and HNF+ LNETs together. Motif analysis (Suppl. Table 2) was conducted using HOMER’s findMotifsGenome looking at all known motifs from the HOMER library, masking repeat sequences (‘-mask’), and comparing against the union set of H3K27ac peaks above (with ‘-bg’). When motif analysis was performed on differential peaks identified from acetylation over the entire peak (DESeq2 analysis between subtypes or between neuroendocrine tumors), we looked for motifs in the entire H3K27ac peak (‘-size given’). When motif analysis was performed on enhancer clusters that were clustered by acetylation of the center 1kbp, we looked for motifs in the center 1kbp (‘-size 1000’). To pick the top 3 motif families (Fig. 1) we picked the 3 most significant motifs, skipping any motif similar to a motif that was already picked.

H3K27ac data of pancreatic and ileal neuroendocrine tumors were obtained from ref. ^7^ and reprocessed as described above. ChIP-seq data of HNF1A, HNF4A and FOXA3 in HepG2 cells were downloaded from ENCODE^15^, accessions ENCSR633HRJ, ENCSR000BLF and ENCSR092OVN. The binding sites of TCF4 in SK-N-SH neuroblastoma cells were downloaded from ENCODE (accession number ENCSR922RFY). Binding sites from ENCODE were IDR thresholded narrow peaks, and bigWig tracks showing signal p-values. The binding sites of ASCL1 in SH-SY5Y neuroblastoma cells were downloaded from ChIP-Atlas^46^ (bigwig files and peak calls with q < 10^−10^), accessions SRX8670115, and SRX8670119^14^. To generate an enhancer heatmap and correlations, HOMER’s annotatePeaks function was used, with 25bp bins within a 5kb window around all peaks except promoter peaks. A single value from each enhancer of the average acetylation at the center 1kb was used to cluster tumors and enhancers. Tumors were clustered by hierarchical clustering of the Spearman correlations of enhancer acetylation by the shortest distance linkage. The enhancers were clustered using k-means clustering and picking the clustering with a minimal error of 100 iterations. Only enhancers with acetylation signals higher than the median were considered when clustering enhancers and generating the heatmap.

RNA-seq reads were mapped to the hg38 reference genome using STAR 2.7.10a (Ref. ^47^) and Ensembl release 99 annotations. Reads were counted using featureCounts, discarding reads with a mapping quality < 10. Density signals were calculated using deeptools’ bamCoverage^40^ with BPM normalization. Gene set enrichment of genes near differential H3K27ac peaks was performed by associating each enhancer to the nearest TSS, if the distance was < 100kbp. Gene set enrichment analysis was calculated by Fisher exact test using HOMER’s findGO script compared to the MSigDB database^48^. Enrichment was compared against the relevant background list using the -bg parameter of findGO: for RNAseq all genes with > 0.5 TPM in at least one sample, and for genes near differential H3K27ac peaks, all genes near any peak (associating nearest gene up to < 100kbp similarly). Relative enrichment score was calculated as follows: (number of target genes in term * total number of genes)/(total number of target genes * number of genes in term), as suggested in Ref. ^49^.

## Acknowledgments

This study was supported by the Neuroendocrine Tumor Research Foundation (YD) and the Alon Fellowship of the Israeli Council for Higher Education (YD). We thank Yinon Ben-Neriah (HUJI) for providing us with the A51 inhibitor and for helpful discussions. We thank Michal Rabani and Rami Aqeilan (HUJI) for critical reading of the manuscript.

## Supplemental materials

**Supplemental Table 1**: Cohort of tumor samples used in this study.

**Supplemental Table 2**: Known motifs enriched in: (1) shared H3K27ac peaks across neuroendocrine tumors; (2) LNET specific peaks; (3) peaks more acetylated in proneural LNETs compared to HNF+ LNETs; (4) peaks more acetylated in HNF+ LNETs compared to proneural LNETs, and; (5) peaks more acetylated in luminal LNETs compared to other LNETs.

**Supplemental Table 3**: MSigDB gene sets enriched (FDR < 0.01) in: (1) genes near H3K27ac peaks more acetylated in proneural LNETs compared to HNF+ LNETs; (2) genes near H3K27ac peaks more acetylated in HNF+ LNETs compared to proneural LNETs; (3) genes near H3K27ac peaks more acetylated in Luminal LNETs; (4) genes near H3K27ac peaks less acetylated in Luminal LNETs; (5) genes downregulated in H835 cells after A51 treatment; (6) genes upregulated in H835 cells after A51 treatment; (7) genes downregulated in H727 cells after A51 treatment; (8) genes upregulated in H727 cells after A51 treatment.

## Notes

### Competing Interest Statement

The authors have declared no competing interest.

### Summary of Updates

Characterized 5 additional tumors and deepened the analysis.

