## Supplemental Figures for "Enhancer landscape of lung neuroendocrine tumors reveals regulatory and developmental signatures with potential theranostic implications"

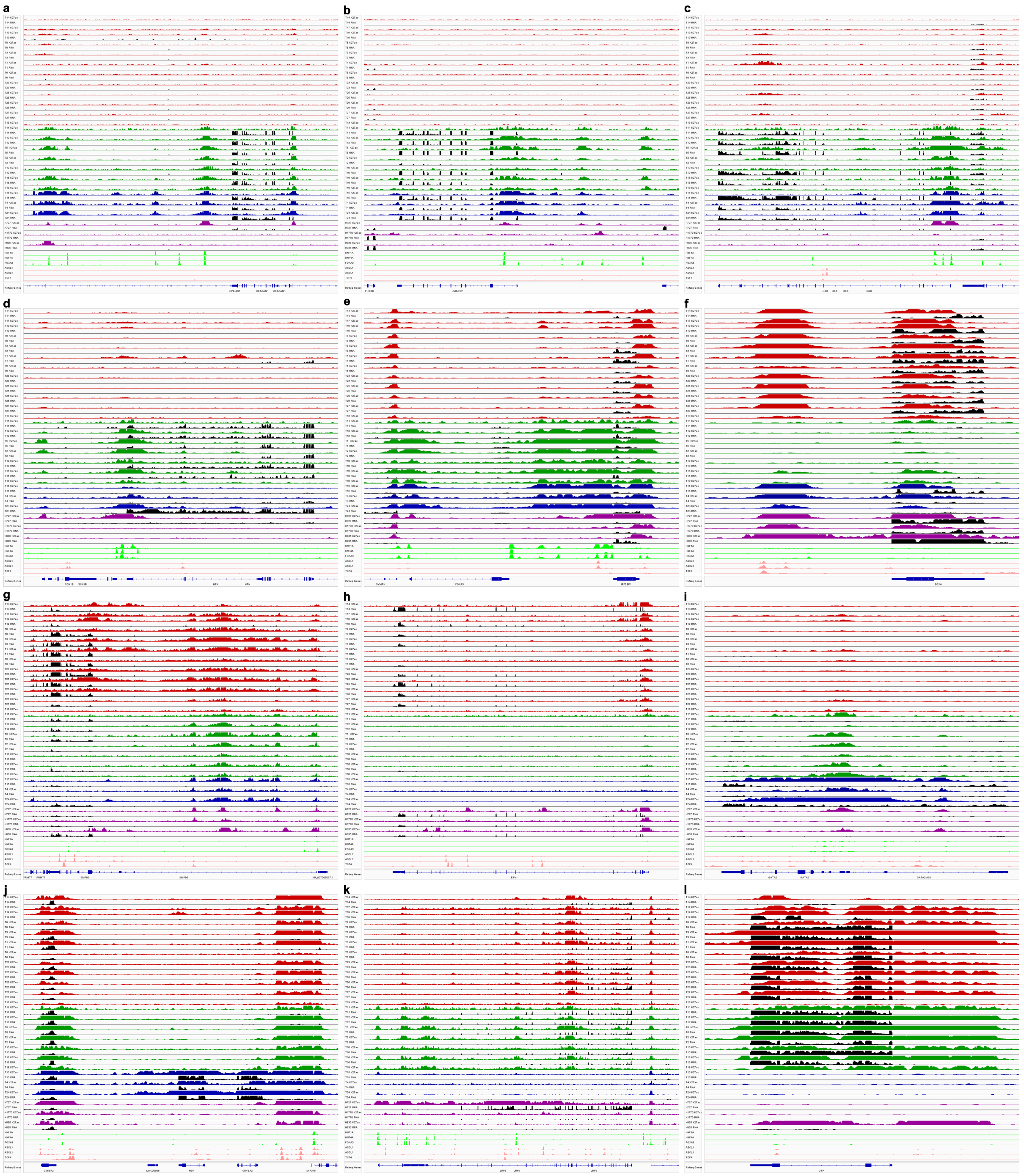

**Supp. Fig. 1: (a-g)** Genomic view of the **(a)** CEACAM1, **(b)** HMGCS2, **(c)** HGD, **(d)** HPN, **(e)** FOXA3, **(f)** SOX4, **(g)** SMPD3, **(h)** ETV1, **(i)** GATA2, **(j)** FEV, **(k)** LRP5 and **(l)** OTP loci, showing RNA signal (black) and H3K27ac signals of proneural (red), HNF+ (green) and luminal (blue) lung neuroendocrine tumors, as well as the NCI-H727, NCI-H1770 and NCI-H835 lung neuroendocrine neoplasms cell lines (purple). Hepatic factors (HNF1A, HNF4A and FOXA3) ChIP-seq tracks in HepG2 cells are shown in light green and proneural transcription factors (ASCL1 and TCF4) ChIP-seq tracks in neuroblastoma cells in light red. RNA-seq signals are normalized to 5 bins per million mapped reads (BPM). H3K27ac signals are scaled by promoter-based DESeq2 normalization (see Methods).

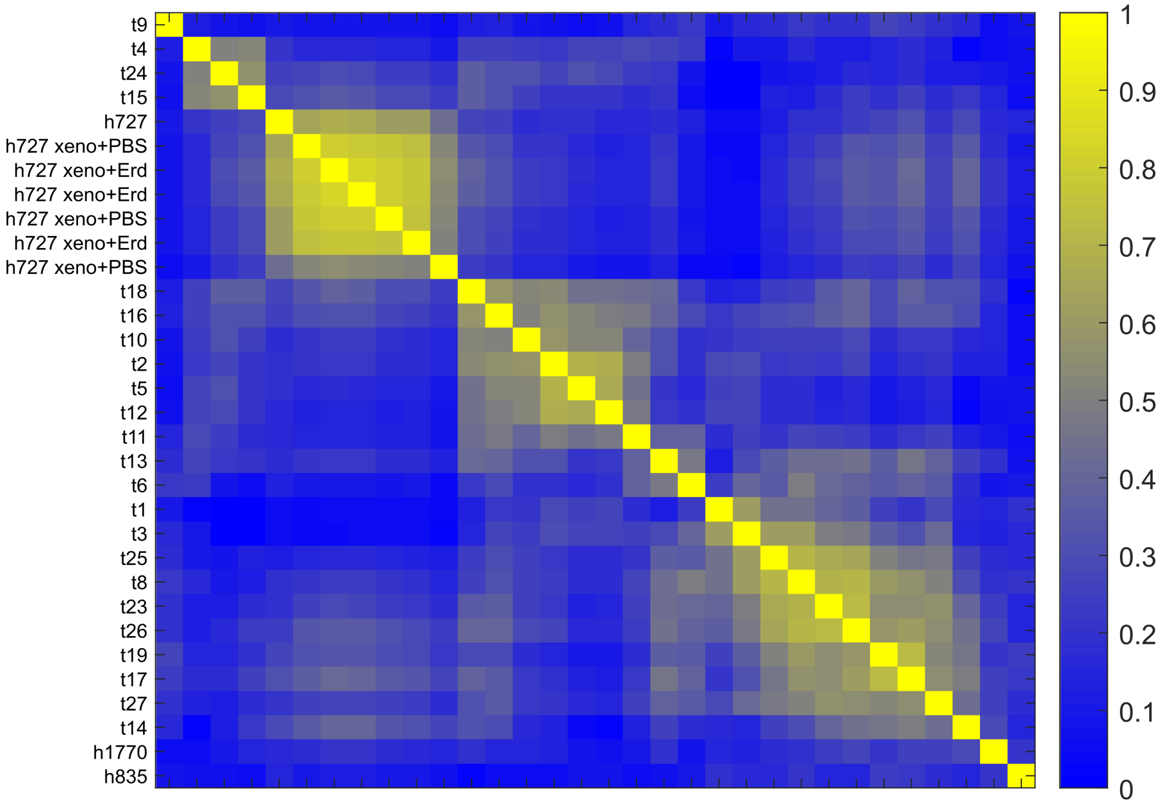


**Supp. Fig. 2:** Pairwise Spearman correlations of H3K27ac signals at distal regulatory elements in lung neuroendocrine tumors, the NCI-H727, NCI-H1770 and NCI-H835 lung neuroendocrine neoplasms cell lines, and six mouse xenograft models generated from NCI-H727 cells – 3 treated with Erdafitinib (12.5mg/kg) and 3 controls treated with PBS. Enhancer landscapes of cell line models and cell line derived xenografts are very different than tumors resected from patients. The global enhancer landscape is maintained upon Erdafitinib treatment.

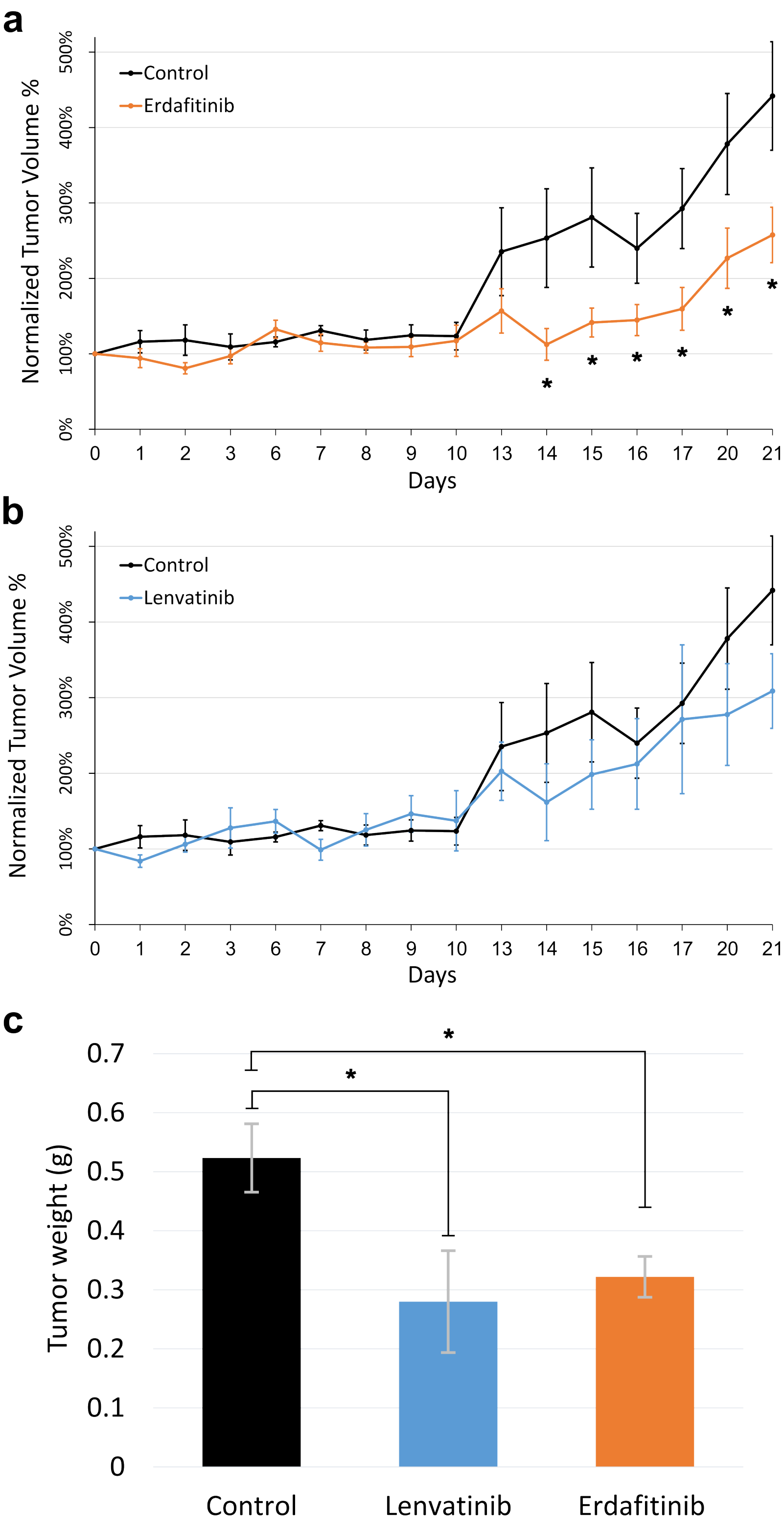


**Supp. Fig. 3: (a)** Normalized tumor volume of NCI-H727 mouse xenograft model treated with Erdafitinib (12.5mg/kg, orange) or 0.5% carboxymethylcellulose control (black). Error bars represent standard error of the mean. P-values are calculated by one tail t-test, * $p < 0.05$. **(b)** Normalized tumor volume of NCI-H727 mouse xenograft model treated with Lenvatinib (5mg/kg, blue) or 0.5% carboxymethylcellulose control (black). Error bars represent standard error of the mean. **(c)** Average tumor weight of the CMC control group (black), Lenvatinib (5mg/kg, blue) and Erdafitinib treated group (12.5mg/kg, orange) at the end of the experiment after mice were sacrificed. Error bars represent standard error of the mean. P-values are calculated by one tail t-test, * $p < 0.05$.
